# Seminal fluid protein divergence among populations exhibiting postmating prezygotic reproductive isolation

**DOI:** 10.1101/2020.06.08.140020

**Authors:** Martin D. Garlovsky, Caroline Evans, Mathew A. Rosenow, Timothy L. Karr, Rhonda R. Snook

**Author notes:** Department of Biology, Syracuse University, Syracuse NY, USA.

## Abstract

Despite holding a central role for fertilisation success, reproductive traits often show elevated rates of evolution and diversification. The rapid evolution of seminal fluid proteins (Sfps) within populations is predicted to cause mis-signalling between the male ejaculate and female reproductive tract between populations resulting in postmating prezygotic (PMPZ) isolation. Crosses between populations of *Drosophila montana* show PMPZ isolation in the form of reduced fertilisation success in both noncompetitive and competitive contexts. Here we test whether male ejaculate proteins deriving from either the accessory glands or the ejaculatory bulb differ between populations using liquid chromatography tandem mass spectrometry. We find more than 150 differentially abundant proteins between populations which may contribute to PMPZ isolation. These proteins include a number of proteases and peptidases, and several orthologs of *D. melanogaster* Sfps, all known to mediate fertilisation success and which mimic PMPZ isolation phenotypes. Males of one population typically produced greater quantities of Sfps and the strongest PMPZ isolation occurs in this direction. The accessory glands and ejaculatory bulb have different functions and the ejaculatory bulb contributes more to population differences than the accessory glands. Proteins with a secretory signal, but not Sfps, evolve faster than non-secretory proteins although the conservative criteria used to define Sfps may have impaired the ability to identify rapidly evolving proteins. We take advantage of quantitative proteomics data from three *Drosophila* species to determine shared and unique functional enrichments of Sfps that could be subject to selection between taxa and subsequently mediate PMPZ isolation. Our study provides the first high throughput quantitative proteomic evidence showing divergence of reproductive proteins implicated in the emergence of PMPZ isolation between populations.

**IMPACT SUMMARY:** Identifying traits that prevent successful interbreeding is key to understanding early stages of the formation of new species, or speciation. Reproductive isolation arising prior to and during fertilisation frequently involves differences in how the sexes interact. In internally fertilising taxa, such interactions are mediated between the female reproductive tract where fertilisation occurs and the receipt of the ejaculate necessary for fertilisation. Because ejaculate proteins are at least partially responsible for these interactions, differences in male ejaculate protein composition could negatively impact fertilisation success, generating reproductive isolation. While the biological classes of ejaculate proteins are shared across all animal taxa, proteins that are secreted by males tend to show rapid evolution in gene expression and genetic sequence. Thus, reproductive proteins are suggested as prime targets facilitating reproductive isolation that arises after mating but before fertilisation (PostMating PreZygotic or PMPZ isolation). Most research on PMPZ isolation has focussed on differences between species for which it is not possible to determine the causative and temporal order of early speciation processes. Here, we test whether populations that exhibit few genetic differences but show strong PMPZ isolation also exhibit variation in ejaculate composition using quantitative high throughput proteomic analyses. We find a number of proteins are differentially abundant between populations including several known to impact fertilisation success in other species. We show that secreted proteins are evolving at an elevated rate, implicating their potential role in PMPZ isolation. We test divergence in ejaculate composition between species, finding a core set of functions that were conserved across species which last shared a common ancestor more than 40 million years ago along with species-specific investment. This work highlights the divergent evolution of reproductive proteins which may contribute to barriers between populations within a species early during speciation, extendable to similar analyses in other taxa in the future.

## INTRODUCTION

For internally fertilising taxa the male ejaculate and female reproductive tract must interact during and after mating to ensure optimal fertility (Pitnick et al. 2009, 2020). In polyandrous species, ejaculate x female reproductive tract interactions are subject to rapid coevolution and diversification due to postcopulatory sexual selection (sperm competition and cryptic female choice) and sexually antagonistic coevolution (Birkhead and Pizzari 2002; Sirot et al. 2015; Firman et al. 2017; Meslin et al. 2017). Thus, despite holding a central role for fertilisation, ejaculate and female reproductive tract traits often show elevated rates of molecular and morphological evolution (Rowe et al. 2015; Ahmed-Braimah et al. 2017; Meslin et al. 2017; VanKuren and Long 2018; Simmons and Fitzpatrick 2019; McGeary and Findlay 2020). Divergence between populations in these traits is predicted to result in the early emergence of reproductive isolation that occurs after mating but before fertilisation (postmating prezygotic; PMPZ) (Lande 1981; Gavrilets 2000; Panhuis et al. 2001). Studies have increasingly documented PMPZ isolation, including prior to any postzygotic isolation (Howard et al. 2009; Bono et al. 2011; Sagga and Civetta 2011; Manier et al. 2013; Jennings et al. 2014; Cramer et al. 2016; Devigili et al. 2018; Garlovsky and Snook 2018; Turissini et al. 2018). In the *Drosophila melanogaster* subgroup, PMPZ isolation accumulates quickly measured by relative rates of evolution of different types of reproductive isolating mechanisms and suggests that PMPZ isolation is important in promoting new species and maintaining species barriers (Turissini et al. 2018). Despite the increasing recognition of the importance of PMPZ isolation, there is little understanding of the molecular basis of ejaculate x female reproductive tract interactions that may generate such barriers (McDonough et al. 2016).

The ejaculate consists of sperm and male seminal fluid proteins (Sfps) that impact fertilisation (e.g., Avila et al. 2011; South and Lewis 2011). Most Sfps are products of male secretory glands (e.g. in mammals, the prostate gland and seminal vesicles; in arthropods, accessory glands and ejaculate ducts/bulb; for review, see Sirot et al. 2015; Fig. S1). Different secretory organs contribute distinct sets of proteins to the ejaculate allowing increased complexity and modulation or tailoring of the ejaculate (Bayram et al. 2019). The majority of work on insect Sfp evolution has been done on *Drosophila melanogaster* with over 200 Sfps identified (Mueller et al. 2005; Findlay et al. 2008, 2009). However, many of the biochemical classes of Sfps are similar across animals; for example, proteases and protease inhibitors, and those with antimicrobial/immune related functions (Avila et al. 2011; Sirot et al. 2015). Despite conserved protein classes observed in the seminal fluid of all animals, a large fraction of Sfps show rapid molecular evolution and, therefore, even Sfps of the same classes in different species are not orthologous (Avila et al. 2011; Perry et al. 2013; Sirot et al. 2015). Functional confirmation of Sfps, performed mostly in *D. melanogaster*, indicate they aid in sperm transfer and storage, influence the outcome of sperm competition, and/or alter female physiology, behaviour and reproductive tract morphology after mating (Ravi Ram and Wolfner 2007; Wong et al. 2008; Avila and Wolfner 2009; Holman 2009; Wigby et al. 2009; Wolfner 2009; Avila et al. 2011; Fedorka et al. 2011; Mattei et al. 2015). Moreover, some Sfps elicit PMPZ-like phenotypes when genetically manipulated (Ravi Ram and Wolfner 2007; LaFlamme et al. 2012). Differences in Sfp expression between species or abnormal gene expression profiles in the female reproductive tract after mating with hetero- vs. con- specific males are associated with PMPZ isolation (Bono et al. 2011; Ahmed-Braimah et al. 2017). These shared and divergent patterns support Sfps as putative causative agents of PMPZ isolation.

However, while studies showing divergence in gene expression are associated with disrupted ejaculate x female reproductive tract interactions, changes in gene expression may not correlate with changes in protein abundance (Wang et al. 2019), where the molecular interactions causing PMPZ isolation take place. Divergence in protein identity or abundance between taxa could disrupt ejaculate x female reproductive tract interactions leading to PMPZ isolation (Goenaga et al. 2015). High-throughput proteomics using liquid chromatography tandem mass spectrometry (LC-MS/MS) has revolutionised identification and quantification of Sfps, revealing that the male ejaculate often contains hundreds of unique proteins (McDonough et al. 2016; Bayram et al. 2019; Karr 2019; Rowe et al. 2019; Whittington et al. 2019). Using LC-MS/MS combined with genomics, Sfps can be predicted by identifying ejaculate proteins with a signal peptide sequence, sometimes called the “secretome”, and those secretome proteins that have an extracellular signal sequence, sometimes called the “exoproteome” (Avila et al. 2011; Ahmed-Braimah et al. 2017; Bayram et al. 2019; Karr et al. 2019; Sepil et al. 2019). Molecular evolution analyses show that proteins with secretory signal and/or are secreted evolve faster than non-secreted ejaculate proteins (Mueller et al. 2005; Wagstaff and Begun 2005; Ramm et al. 2009; Bono et al. 2015; Tsuda et al. 2015; Ahmed-Braimah et al. 2017; Karr et al. 2019)

While these results suggest Sfps are prime candidates for generating PMPZ isolation, no study using high throughput quantitative proteomics has tested the prediction that ejaculate composition will vary between populations exhibiting PMPZ isolation, and that these proteins will more rapidly evolve. Similarly, while previous work has suggested that the different ejaculate secretory organs may perform different functions (Bayram et al. 2019), their potential contribution to PMPZ isolation has not been examined. Here we use LC-MS/MS on the accessory glands and the ejaculatory bulb/duct, followed by molecular evolutionary rates analysis, to test these predictions using the malt fly, *Drosophila montana*. We have focused on two populations (Crested Butte, Colorado; Vancouver, Canada) and found that reciprocal crosses between populations result in PMPZ isolation both after a single mating as males transfer sperm to females who store them, but many eggs are not fertilised (Jennings et al. 2014; Garlovsky and Snook 2018), and where within- and between- population males compete for fertilisation (frequently referred to as conspecific sperm precedence) (Garlovsky et al. in review). The similar results suggest a shared mechanism generating PMPZ isolation. Genomic analysis found no fixed SNPs between these populations, likely due to a history of gene flow during divergence (Parker et al. 2018; Garlovsky et al. in review), although genes enriched for reproductive function show divergence (Parker et al. 2018). These results support focussing on Sfps as potential causative agents of PMPZ isolation. We also leverage recent high throughput mass spectrophotometry data on ejaculate composition in two other *Drosophila* species (Karr et al. 2019; Sepil et al. 2019) to provide insights into shared and divergent Sfp functions that may contribute to PMPZ isolation.

## METHODS

### Fly stocks

Adult *Drosophila montana* were collected with malt bait buckets and mouth aspirators in Crested Butte, Colorado, USA (38°49’N, 107°04’W) in 2013 (referred to as Colorado), and Vancouver, British Columbia, Canada (48°55’N, 123°48’W) in 2008 (referred to as Vancouver) (Fig. S1). Stocks were established by combining 20 F3 males and females from 20 isofemale lines (800 flies total per population) and cultured on Lakovaara malt media (Lakovaara 1969) in overlapping generations in constant light at 19°C. Flies were collected within 3 days of eclosion and housed in groups of between 10-20 single sex individuals in food vials until reproductively mature at 21 days old.

### Tissue collection and protein extraction

Twenty-one-day old males were anaesthetised with ether and the accessory glands and ejaculatory duct/bulb separated from nontarget tissues, and from each other. We collected three biological replicates, two of which were separated into technical replicates (Fig. S1). Following protein extraction and purification, we quantified protein concentration to standardize loading 5μg of protein for each sample into the mass spectrometer (see supplementary material; Fig. S2). Samples were reduced with TCEP, alkylated by addition of MMTS, and digested with trypsin, followed by drying to completion using vacuum centrifugation. Samples were resuspended in 20μl 3% v/v acetonitrile, 0.1% v/v trifluoroacetic acid prior to LC-MS/MS analysis.

### LC-MS/MS analysis

Detailed description of LC-MS/MS data acquisition and processing can be found in the supplementary material.

LC-MS/MS was performed by nano-flow liquid chromatography (U3000 RSLCnano, Thermo Fisher™) coupled to a hybrid quadrupole-orbitrap mass spectrometer (QExactive HF, Thermo Scientific™). Peptides were separated on an Easy-Spray C18 column (75 μm × 50 cm) using a 2-step gradient from 97% solvent A (0.1% formic acid in water) to 10% solvent B (0.08% formic acid in 80% acetonitrile) over 5 min then 10% to 50% B over 75 min at 300 nL/min. The full 105-minute MS data dependent acquisition was set up from 375-1500 m/z acquired in the Orbitrap in profile mode, resolution 120,000. Subsequent fragmentation was Top 10 in the HCD cell, with detection of ions in the Orbitrap using centroid mode, resolution 30,000. MS parameters; MS1: Automatic Gain Control (AGC) target 1e6 with a maximum injection time (IT) of 60 ms; MS2: AGC target 1e5, IT of 60 ms and isolation window 2 Da.

We performed label free quantitative proteomic analysis using MaxQuant to generate relative peptide and protein intensities (Cox et al. 2014; Tyanova et al. 2016) (see supplementary material). For protein identification we matched mass spectra to the *D. montana* predicted proteome (Parker et al. 2018), generated using gene predictions from the Maker2 pipeline (Holt and Yandell 2011) reciprocally blasted against *D. virilis* proteins (Parker et al. 2018).

### Gene Ontology (GO) and functional analysis

We performed network analyses and GO enrichment for Biological Processes (BP), Cellular Components (CC) and Molecular Functions (MF) with the ClueGO plugin (Bindea et al. 2009) for Cytoscape (Shannon et al. 2003) using FlyBase gene numbers (FBgns) for *D. virilis* orthologs of *D. montana* genes retrieved from Parker et al. (2018) or *D. melanogaster* orthologs converted via FlyBase.org. Specific settings for network groups are provided in figure and table legends. For GO enrichment we used right-sided hypergeometric tests with Benjamini-Hochberg multiple test correction.

### Differential abundance analysis between D. montana populations and functional differences between tissues

We performed differential abundance analysis of MaxLFQ ion intensities using the ‘edgeR’ (Robinson et al. 2010) and ‘limma’ (Ritchie et al. 2015) packages in R (v.3.5.1) (R Core Team 2018) (see supplementary material). Proteins were considered differentially abundant based on an adjusted p-value < 0.05. To identify differentially abundant proteins between populations, we analysed the accessory gland proteome and the ejaculatory bulb proteome separately. We only considered proteins that were present in all five replicates of each tissue for both populations. To identify differentially abundant proteins between tissues, we analysed each population separately. Again, we only considered proteins that were present in all five replicates of each population for both tissues. (Table S1).

### Characterising the male seminal fluid proteome across species

We compared Sfp functions for three *Drosophila* species for which proteomic data generated using LC-MS/MS is available for the male accessory gland and ejaculatory duct and bulb tissues: *D. montana* (this study), *D. melanogaster* (Sepil et al. 2019), and *D. pseudoobscura* (Karr et al. 2019). We retrieved FBgns for *D. melanogaster* genes identified by Sepil et al. (2019) and *D. melanogaster* orthologs for *D. pseudoobscura* genes identified by Karr et al. (2019) and downloaded the corresponding canonical protein sequences from uniprot.org. For proteins we identified in our analysis we retrieved *D. montana* protein sequences from Parker et al. (2018). We submitted protein sequences for each species to *SignalP* (Petersen et al. 2011) and *Phobius* (Käll et al. 2004) and combined the resulting lists of proteins containing a signal peptide to generate a list of secretome proteins for each species. For *D. montana* we converted the corresponding *D. virilis* FBgns for each protein to *D. melanogaster* orthologs via FlyBase.org (for 215/245, 88%). To identify Sfps for each species we submitted secretome lists to FlyBase.org to retrieve genes with GO terms containing “extracelluar” (Fig. S3; Table S2). To compare GO enrichment between species we adjusted network settings in ClueGO to reflect the different numbers of proteins identified in each species.

### Evolutionary rates analysis

To obtain sequence divergence estimates for *D. montana* proteins we used a pipeline developed previously (Wright et al. 2015). We obtained protein coding sequences for *D. montana* from Parker et al. (2018) and for *D. pseudoobscura* (r3.04, September 2019) and *D. virilis* (r1.07, August 2019) from FlyBase.org. We identified the longest isoform of each gene for each species and determined orthology with reciprocal BLASTN (Altschul et al. 1990), using a minimum percentage identity of 30% and an E-value cut-off of 1×10^−10^. We then identified reciprocal one-to-one orthologs across all three species using the highest BLAST score. We identified open reading frames using BLASTx and aligned orthologs using PRANK (Löytynoja and Goldman 2010). We calculated the ratio of non-synonymous (dN) to synonymous (dS) nucleotide substitutions, omega (ω), using the CODEML package in PAML (Yang 2007) (one-ratio estimates, model 0) with an unrooted phylogeny. Results were filtered to exclude orthologs with branch-specific dS ≥ 2 (due to potential mutational saturation) or where S*dS ≤ 1.

We then tested for differences in evolutionary rates between sets of proteins we identified in our LC-MS/MS analysis. We relaxed filtering criteria so that a protein need only be identified in a single replicate in a single population or tissue, but still had to be identified by two or more unique peptides. After filtering, we obtained ω values for 757/1474 (51%) proteins with a reciprocal one-to-one ortholog. We categorised genes as having higher abundance in either the accessory gland proteome or ejaculatory bulb proteome as those proteins showing concordant differential abundance between tissues across populations (Fig. 3). We classified proteins as belonging to the secretome based on presence of a signal peptide and as Sfps as secretome proteins with extracellular annotation (see above) plus *D. melanogaster* Sfps. We classified the remainder of proteins excluding those with higher abundance in the accessory gland proteome, ejaculatory bulb proteome, secretome, or Sfps, as background tissue. Each class consisted of a unique set of proteins, such that accessory gland proteins did not include ejaculatory bulb proteins, secretome, Sfps, or remaining proteins; secretome proteins did not include Sfps, etc. We tested for differences in evolutionary rates using the Kruskal-Wallis rank sum test followed by pairwise Wilcox rank sum tests corrected for multiple testing using the Benjamini-Hochberg method.

## RESULTS

### The *D. montana* accessory gland proteome and ejaculatory bulb proteome

We identified 1711 proteins, of which 1474 (86%) were identified by two or more unique peptides. The majority of proteins (1013/1474; 69%) were shared across male secretory tissues, while 138 (9%) and 323 (22%) proteins were unique to the accessory glands and ejaculatory bulb, respectively (Fig. S4a). Proteins identified only in the accessory gland proteome showed a 3.2-fold lower mean abundance compared to the remainder of proteins whereas proteins identified only in the ejaculatory bulb proteome showed a 14.9-fold reduction. These proteins likely represent missed rather than truly unique proteins and are not considered further. We identified 79 *D. montana* Sfps, consisting of 38 orthologs of *D. melanogaster* Sfps identified by converting *D. virilis* FBgns on FlyBase.org, plus 55 secretome proteins with extracellular annotations identified by 2 or more unique peptides (14 of which overlapped) (Fig. S4a; Table S3) (Mueller et al. 2005; Findlay et al. 2008, 2009). A multidimensional scaling (MDS) plot of normalised intensities using all proteins (n = 1474) showed a clear separation of samples by tissue type (dimension 1), and separation by population (dimension 2) with clear separation of populations for the accessory gland proteome and marginal overlap between populations in the ejaculatory bulb proteome (Fig. 1).

**Figure 1.**
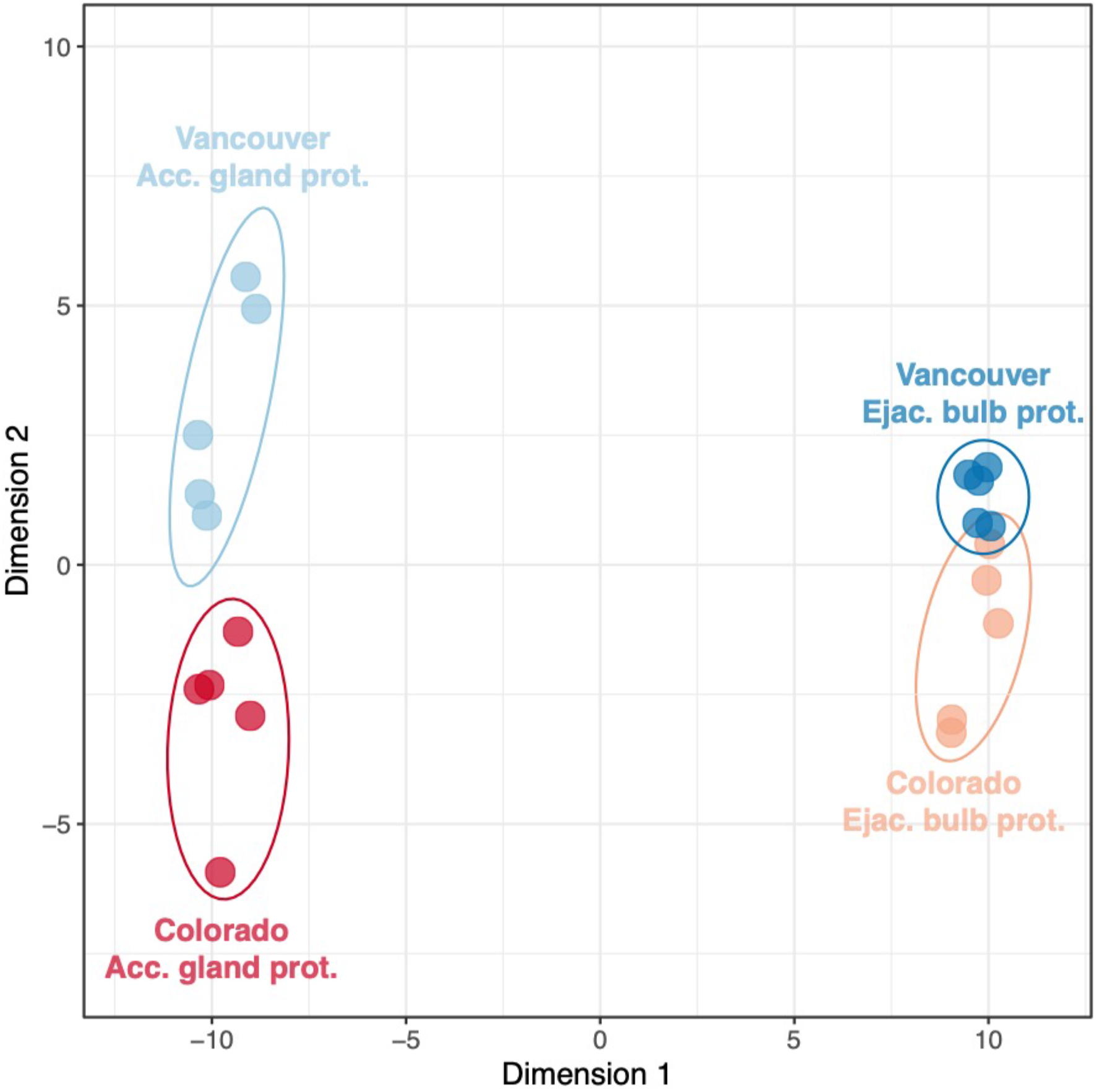
Multidimensional scaling (MDS) plot of normalised intensities for proteins identified by two or more unique peptides (n = 1474) in each replicate (points). Dimension 1 separates the two tissue types (accessory glands and ejaculatory bulbs). Dimension 2 separates the two populations (Colorado and Vancouver).

### Differential abundance of reproductive proteins between populations

The majority of proteins were identified in both populations (1322/1474; 90%), while 45 (3%) and 107 (7%) were only identified in Colorado, and Vancouver, respectively (Fig. S4b). Proteins only identified in one population showed a 263-fold, and 171-fold, lower mean abundance compared to the rest of proteins in Colorado, and Vancouver, respectively. As above, these low abundance proteins are not considered further. For shared proteins, we then tested for differential abundance. We identified 154 (out of 725) differentially abundant proteins produced in the accessory glands between populations (Fig. 2a), including nine *D. melanogaster* Sfps (Table 1). We identified 244 (out of 929) differentially abundant proteins produced in the ejaculatory bulbs (Fig. 2b). Again, this included nine *D. melanogaster* Sfps, two of which overlapped with those identified in the accessory gland proteome (Table 1). In the accessory gland proteome, Sfps and proteins with a predicted secretory signal were not overrepresented in the cohort of proteins showing differential abundance (Chi-squared test, Χ^2^ = 1.57, df = 2, p = 0.456) but were overrepresented in the cohort of differentially abundant proteins in the ejaculatory bulb proteome (Χ^2^ = 44.56, df = 2, p < 0.001; Fig. S5). Out of 45 proteins that were differentially abundant between populations in both male reproductive tissues, 36 showed higher abundance in one population in both tissues (Fig. S6). Significantly enriched gene ontology (GO) categories for proteins showing differential abundance between populations are in Tables S4-S6.

**Figure 2.**
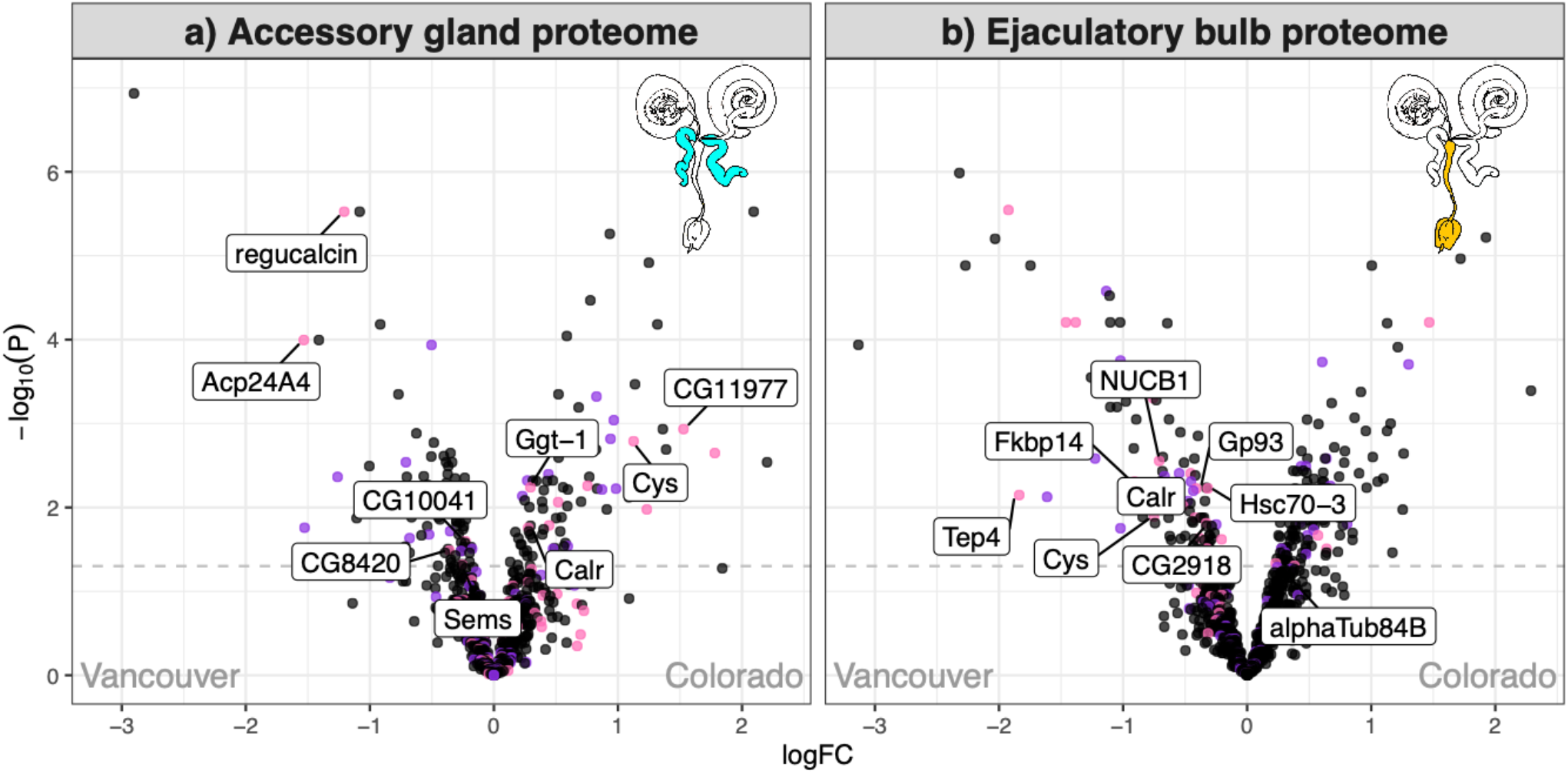
Differentially abundant proteins between Colorado and Vancouver in a) the accessory glands (n = 725) and b) the ejaculatory bulbs (n = 929). Secretome proteins are shown in purple and Sfps in pink. Significantly differentially abundant proteins with a known Sfp ortholog in *D. melanogaster* are labelled.

**Table 1.** Differentially abundant proteins between *D. montana* populations with a known *D. melanogaster* seminal fluid protein (Sfp) ortholog. Gene names were retrieved from FlyBase.org using the corresponding *D. virilis* FBgns for *D. montana* proteins we identified via LC-MS/MS. The population for which each protein showed higher abundance is given. Proteins found in both tissue comparisons indicate in which population there was higher abundance. Abbreviations: Acgs, accessory glands; Ebs, ejaculatory bulbs.

| Tissue comparison | Gene name | Higher abundance |
| --- | --- | --- |
| Accessory glands | CG11977 | Colorado |
|  | gamma-glutamyl transpeptidase | Colorado |
|  | Acp24A4 | Vancouver |
|  | CG10041 | Vancouver |
|  | CG8420 | Vancouver |
|  | Regucalcin | Vancouver |
|  | Seminase | Vancouver |
| Ejaculatory bulbs | alpha-Tubulin at 84B | Colorado |
|  | CG2918 | Vancouver |
|  | FK506-binding protein 14 | Vancouver |
|  | Glycoprotein 93 | Vancouver |
|  | Heat shock 70-kDa protein cognate 3 | Vancouver |
|  | NUCB1 | Vancouver |
|  | Thioester-containing protein 4 | Vancouver |
| Both | Calreticulin | Colorado<sup>Acgs</sup>/Vancouver<sup>Ebs</sup> |
|  | Cystatin-like | Vancouver<sup>Acgs</sup>/Colorado<sup>Ebs</sup> |

### The accessory gland and ejaculatory bulb proteomes differ in function

To test whether the accessory glands and ejaculatory bulb provide different functions we performed differential abundance analysis between tissues for Colorado and Vancouver separately. We found 524 (out of 652) differentially abundant proteins between tissues in Colorado. Similarly, in Vancouver we found 557 (out of 676) differentially abundant proteins. The majority of these proteins were found in both populations (609 proteins). To identify consistently differentially abundant proteins between tissues, we compared the log2-fold change in abundance in each population of these 609 proteins. Proteins with higher abundance in the accessory gland proteome or ejaculatory bulb proteome in Colorado generally also showed higher abundance in Vancouver (Spearman’s rank correlation, ρ = 0.945, p < 0.001, n = 609) (Fig. 3a). GO analyses identified both tissues as having functions expected for highly metabolically active secretory organs (Table S7). Different GO terms were enriched in each tissue highlighting that the two secretory organs provide distinct reproductive functions to the ejaculate (Fig. 3b; Table S7).

**Figure 3.**
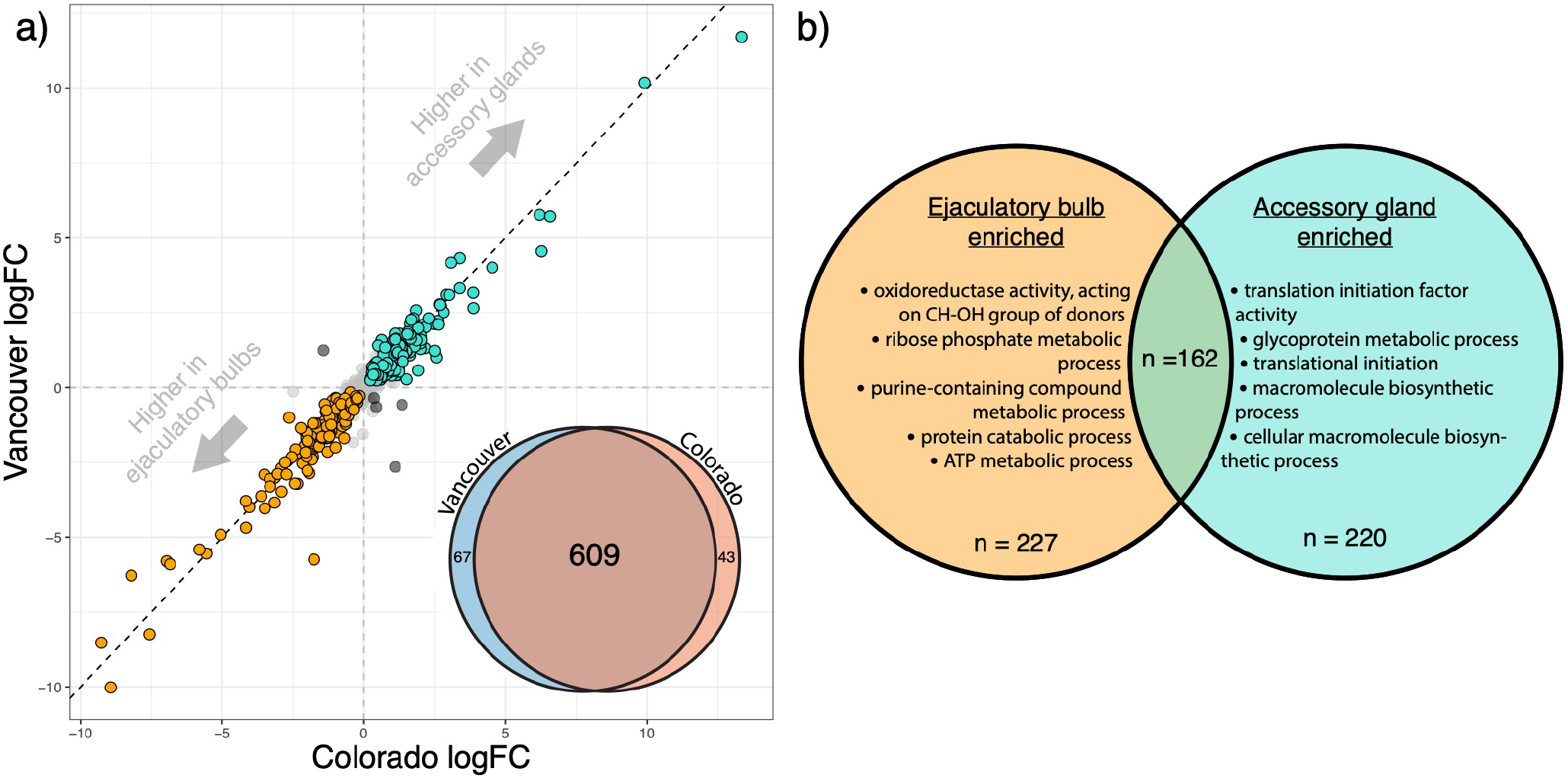
Populations show consistent differential abundance between secretory organs. a) Concordance between populations in log2-fold change in abundance of proteins found in either the accessory gland proteome or ejaculatory bulb proteome. Positive values indicate proteins with higher abundance in the accessory glands in both populations (top right), negative values indicate proteins with higher abundance in the ejaculatory bulb (bottom left). Proteins are coloured based on whether they showed a concordant pattern of significantly higher abundance in the accessory gland proteome (turquoise, n = 220), the ejaculatory bulb proteome (orange, n = 227), were discordant (black, n = 5), or were not significantly differentially abundant between tissues (grey). Dashed black line shows 1:1. Inset: Venn Diagram showing numbers of proteins included in separate differential abundance analysis between tissues in each population and overlap. b) Top 5 significantly enriched Biological Process GO terms ranked by percent identity of proteins to each tissue (see Table S7 for full list).

### Evolutionary rates analysis

We tested whether genes with higher protein abundance in either the accessory gland proteome, ejaculatory bulb proteome (excluding the secretome and Sfps), secretome (excluding Sfps), or Sfps (secretome proteins with extracellular annotation), were evolving at different rates compared to background proteins, defined as those proteins that do not differ in protein abundance between the accessory glands and ejaculatory bulb and excluding secretome and Sfps. There was a significant difference between protein groups in evolutionary rates (Kruskal-Wallis test, Χ^2^ = 40.51, df = 4, p < 0.001) (Fig. 4). The secretome and Sfps were evolving at similar rates (pairwise Wilcox rank sum test with Benjamini-Hochberg adjustment, p = 0.668). The secretome was evolving faster than proteins with higher abundance in the accessory gland proteome, ejaculatory bulb proteome, or background (all p < 0.007). Sfps were also evolving faster than proteins with higher abundance in the accessory gland proteome (p = 0.001) and ejaculatory bulb proteome (p = 0.007) but at a similar rate to background (p = 0.171). Proteins with higher abundance in the accessory gland proteome and ejaculatory bulb proteome were evolving at similar rates (p = 0.089), and slower than the remaining background proteome (accessory gland proteome vs. background, p < 0.001; ejaculatory bulb proteome vs. background, p = 0.011).

**Figure 4.**
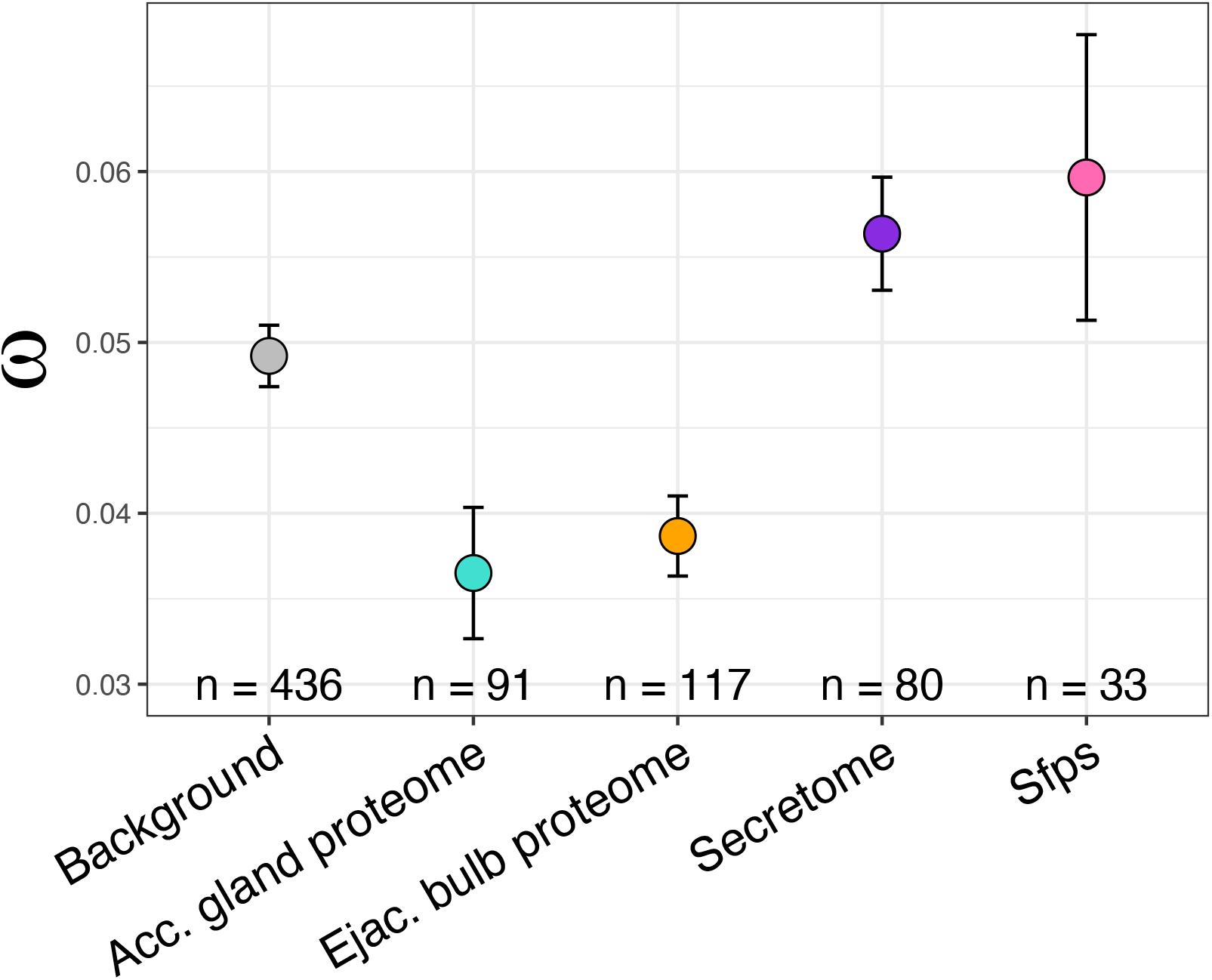
Mean non-synonymous (dN) to synonymous (dS) base substitution rate (ω) estimates (± standard error) for *D. montana* genes identified in our LC-MS/MS analysis with equal abundance in the accessory gland proteome and ejaculatory bulb proteome (‘background’; grey), higher abundance in the accessory gland proteome (turquoise), ejaculatory bulb proteome (orange), or found in the secretome (purple) or Sfps (pink). See Fig. S7 for separate dN and dS plots.

### Comparison of male Sfps across species

We identified 61 Sfps (secretome proteins with extracellular annotations) for *D. montana*, 249 Sfps for *D. melanogaster*, and 131 Sfps for *D. pseudoobscura* (Fig. S3; Table S2). Comparing functional enrichment of Sfps across species identified a number of shared and unique GO categories. Shared Biological Processes included chitin catabolic process, innate immune response, cell-substrate adhesion, and regulation of peptidase activity (Fig. 5; see Table S8 for CC and MF terms). Uniquely enriched BP functions included regulation of secondary metabolic process (*D. montana*); postmating regulation of female receptivity (*D. melanogaster*) and amino glycan catabolic processes (*D. pseudoobscura*) (Fig. 5; see Table S8 for CC and MF terms).

**Figure 5.**
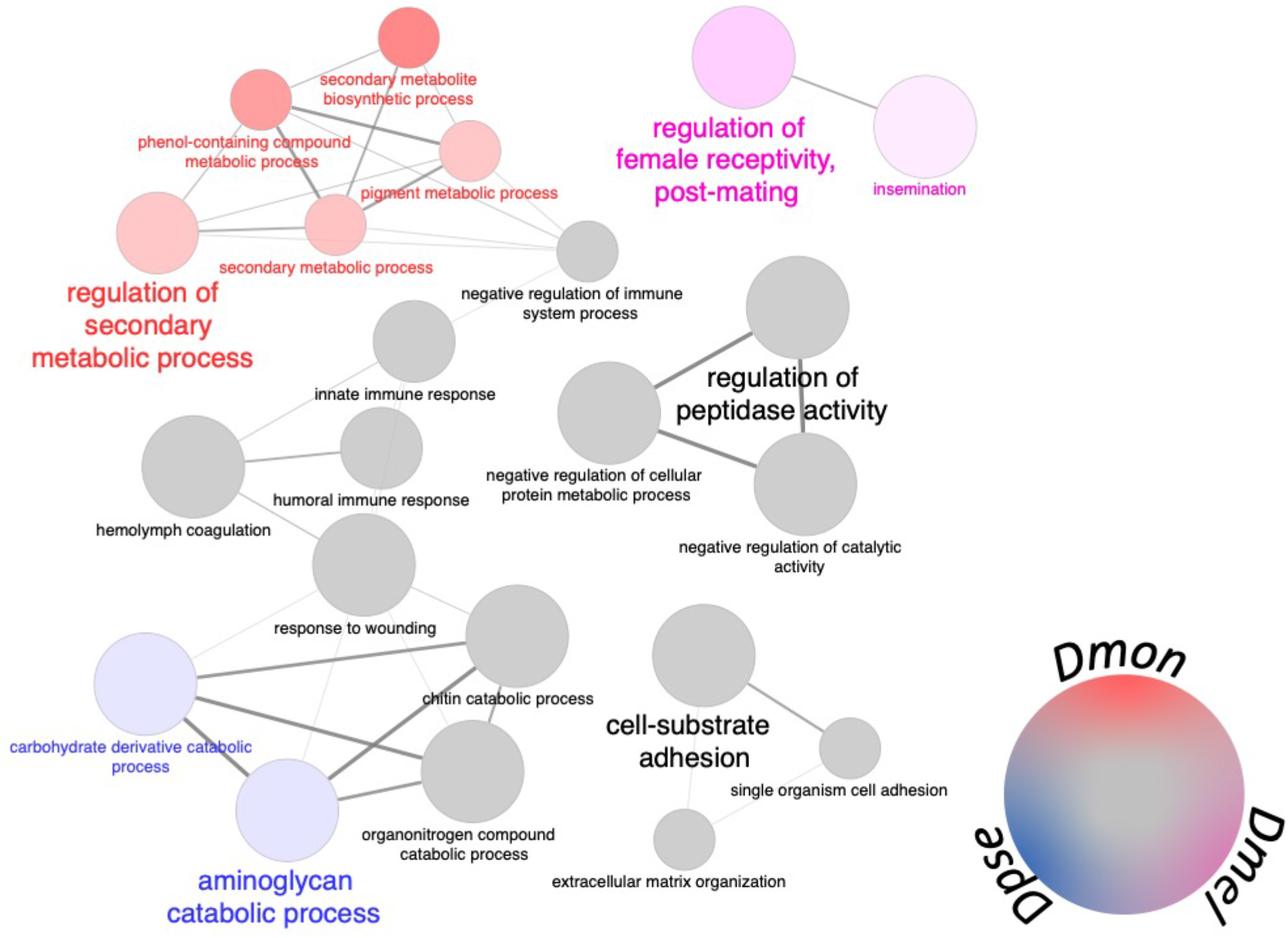
Seminal fluid protein (Sfps) comparisons for GO BP terms across species. Circle size is associated with level of significance with increasing size indicating increasing significance. Node colour indicates proportion of genes from each species associated with a term: *D. montana* (red. Dmon), *D. melanogaster* (pink; Dmel), and *D. pseudoobscura* (blue; Dpse), shared terms are shown in grey. Min. GO level = 3, max. GO level = 8. Number of genes/% genes per group: *D. montana* 3/3%, *D. pseudoobscura* 6/6%, *D. melanogaster* 12/12%. Percentage significance = 55%, kappa-score threshold = 0.25.

## DISCUSSION

The molecular basis of mechanisms underlying PMPZ isolation are poorly understood. Seminal fluid proteins are likely to contribute to PMPZ isolation due to their effects on sperm use, fertilisation success, and rapid divergent evolution. We used quantitative proteomics to identify proteins produced in the accessory glands and ejaculatory duct and bulb in populations exhibiting PMPZ isolation and found a number of differentially abundant proteins between populations including several orthologs of *D. melanogaster* Sfps. The accessory glands and ejaculatory bulb showed different functions and there were more differentially abundant proteins found in the ejaculatory bulb than the accessory glands. For proteins found in both populations, but in separate tissues, there was strong concordance in abundance between populations. We found secretome proteins evolved at a faster rate than non-secretome proteins, both those differentially abundant between male secretory organs and those showing similar abundance between male tissues. Sfps did not show elevated rates of molecular evolution, likely because identification by extracellular annotation required orthology in *D. melanogaster*. Despite shared Sfps, and a core set of shared Sfp biological processes across three *Drosophila* species, there was species-specific enrichment of Sfp function.

*D. montana* from Colorado and Vancouver show low genome-wide divergence and a history of gene flow (Parker et al. 2018; Garlovsky et al. in review), yet show enrichment of genes with reproductive function that are divergent between populations (Parker et al. 2018). Crosses between Colorado and Vancouver show reduced fertilisation success after a single mating (Jennings et al. 2014; Garlovsky and Snook 2018) and exhibit conpopulation sperm precedence (Garlovsky et al. in review). Between different *Drosophila* species, mechanisms causing PMPZ isolation include abnormal sperm transfer and displacement, or mismatches between sperm length and female tract morphology (Price et al. 2001; Manier et al. 2013). Females receiving a foreign ejaculate comprising an abnormal Sfp complement might also result in mismatched ejaculate × female reproductive tract interactions measured by gene expression differences (Bono et al. 2011; Plakke et al. 2015), although variation between species in ejaculate composition has not been quantified in those studies.

We identified a number of differentially abundant proteins between populations exhibiting PMPZ isolation, including several orthologs of *D. melanogaster* Sfps. Intriguingly, 11 of 14 of these proteins were more abundant in Vancouver males than Colorado males. PMPZ isolation between *D. montana* populations is asymmetric, with matings between Vancouver males and Colorado females having lower fertilisation success compared to the reciprocal cross (Jennings et al. 2014; Garlovsky and Snook 2018). If Vancouver males transfer more of these Sfps to their mates, then the chemical environment in the reproductive tract of Colorado females may be mismatched, more so than the reciprocal cross.

Additionally, a number of proteases and peptidases (or inhibitors) were differentially abundant between populations. Proteases and peptidases are central to reproduction across taxa, regulating proteolytic activity and initiating cascades of interactions among downstream proteins (LaFlamme et al. 2012; LaFlamme and Wolfner 2013; Plakke et al. 2015, 2019; Bayram et al. 2017, 2019). Divergence in proteases has been implicated in PMPZ isolation between other insect species in both the male ejaculate and female reproductive tract secretions (Kelleher et al. 2007; Kelleher and Pennington 2009; Marshall et al. 2009, 2011; Meslin et al. 2017; Al-Wathiqui et al. 2018; Plakke et al. 2019).

In *D. montana* females receive and store motile sperm from incompatible males, but fertilisation success is reduced (Jennings et al. 2014). Failure to either properly orient sperm in storage (Manier et al. 2013), release sperm from storage, or have sperm release coincide with ovulation (Mattei et al. 2015) could explain PMPZ in this system (Jennings et al. 2014). Some notable differentially abundant Sfps and proteases we identified, and their potential relationship to PMPZ isolation in *D. montana* are seminase, γ;-glutamyl transpeptidase, and regucalcin. Seminase (CG10586), is a serine protease and a member of the Sex Peptide (SP) network (Singh et al. 2018). Seminase acts early in the SP network and is required to process other Sfps in the mated female essential for proper sperm storage (Acp36DE) and ovulation (ovulin) (LaFlamme et al. 2012; Singh et al. 2018). RNAi knockdown of seminase in male *D. melanogaster* results in failure of mated females to release sperm from the seminal receptacle (LaFlamme et al. 2012). γ;-glutamyl transpeptidase (CG6461) functions to maintain a protective redox environment for sperm (Walker et al. 2006). Mismatches between the male ejaculate and the redox environment of the female reproductive tract in which sperm are stored could reduce fertilisation success as sperm subject to increased oxidative stress are less fertilisation competent (Reinhardt and Ribou 2013). Regucalcin (CG1803), a Ca^+2^ binding protein, may also play an anti-oxidative role and, in mammals, is hypothesized to have an anti-capacitation role for sperm (Pillai et al. 2017). One aspect of capacitation, hyperactivation, increases sperm motility which is important for sperm storage in *Drosophila* (Köttgen et al. 2011). Sperm motility behaviour and how this may affect release from storage is unknown. Regucalcin gene expression varies between *D. montana* populations and has been suggested as a cold tolerance gene in diapausing females (Vesala et al. 2012) although it’s expression in males has not been studied. These examples provide strong candidates for eliciting PMPZ isolation and will be subject to future studies, for instance using CRISPR/Cas9 gene editing, to further understand the molecular interactions causing PMPZ isolation in *D. montana*.

Reproductive proteins are predicted to evolve rapidly, driven by postcopulatory sexual selection and sexual conflict (Sirot et al. 2015; Ahmed-Braimah et al. 2017; Firman et al. 2017; Meslin et al. 2017). We found that proteins showing secretory signals (secretome) evolve faster than proteins without this signal. Sfps and secretome proteins were evolving at a similar rate. However, Sfps were not evolving faster than proteins with similar abundance between male tissues, possibly due to showing greater variation than other categories despite having a higher mean rate. The requirement to have extracelluar annotation determined from work in *D. melanogaster* limits our ability to identify rapidly evolving Sfps in *D. montana*. Thus, the putative number of Sfps in *D. montana*, 79, is a conservative estimate. Isotopically labelling males to identify proteins transferred to females increased the number of identified *D. melanogaster* Sfps (Findlay et al. 2008, 2009). Future work on *D. montana* can use this technique to identify additional Sfps.

One goal of this work was to assess whether the different male reproductive secretory organs had different functions, which would not be possible using the heavy labelling technique. Our work provides one of the first proteomic descriptions of both major Sfp secretory organs in *Drosophila*. Previous research in seed beetles has shown that division of labour across secretory organs enables increased complexity to the ejaculate and potential for ejaculate tailoring (Bayram et al. 2019). Most proteins we identified were found in both tissues but showed higher abundance in either the accessory glands or ejaculatory duct and bulb, suggesting these organs provide different functions to the ejaculate. The accessory gland proteome was enriched for translation and biosynthetic processes, whereas the ejaculatory bulb proteome showed enrichment for mainly metabolic processes. We found secretome proteins and Sfps were significantly overrepresented in the set of differentially abundant proteins in the ejaculatory bulb proteome but not the accessory gland proteome, suggesting the two male secretory organs may contribute differently to PMPZ isolation. Most past work on Sfp evolution has focused on the accessory glands, which could skew understanding of not only molecules involved in reproduction but those reproductive molecules that may elicit PMPZ isolation.

We took advantage of recent accessory gland proteomes for three *Drosophila* species generated using high throughput LC-MS/MS to characterise shared and enriched functions of Sfps between species. Using the same identification criteria for all species (secretory sequence and extracellular annotation), we identified a set of shared GO categories between species that last shared a common ancestor 40 million years ago. This core set included immune-related genes, which are associated with sexual conflict in *D. melanogaster* (Innocenti and Morrow 2009). We also found species-specific GO-enrichment of Sfps, suggesting divergence in how they contribute to male ejaculate function between species. Differences may reflect how selection has targeted particular ejaculate traits in different mating systems (Markow 2002). Differences will also reflect the use of *D. melanogaster* as the reference for GO annotation. For instance, *D. melanogaster* showed enrichment for reproductive genes but Sfps in the other species clearly have reproductive functions. It is likely that reproductive genes which experience strong divergent selection may no longer resemble *D. melanogaster* genes. Our work offers a first insight into the proteomic composition of male ejaculate characteristics across species. As understanding of the molecular interactions between the sexes matures, it will be important to determine whether shared or divergent functions between species are more likely to contribute to PMPZ isolation and when during speciation such divergence occurs. Are shared functions more likely to diverge within populations early during speciation or are Sfps that already show some species specificity more likely to contribute to early PMPZ isolation?

Here we have tested whether reproductive proteins show differential abundance between populations that exhibit PMPZ isolation. Our study has focussed on *Drosophila*, a model system for studying the evolution of reproductive processes and the evolution of reproductive isolation in metazoans. However, reproductive processes, classes of reproductive proteins, and the action of PMPZ isolation across animals show similarities. For example, differentially abundant proteins between *D. montana* populations we found included a number of proteases or peptidases which are common and important mediators of reproductive processes in all animals. Differentially abundant proteins also included several orthologs of *D. melanogaster* Sfps with functions that may be similar to altered reproductive processes generating PMPZ isolation in *D. montana*, such as noncompetitive gametic isolation and conspecific sperm precedence (Jennings et al., 2014; Garlovsky et al. in review). These reproductive isolating mechanisms are found in many other metazoan taxa (for brief review, see Turissini et al. 2018). We also showed secretome proteins are evolving faster than other proteins found in the accessory glands or ejaculatory duct and bulb, and at the same rate as Sfps. Such rapid evolution is frequently attributed to sexual selection and sexual conflict, and these dynamic processes may contribute to speciation (Gavrilets 2000; Panhuis et al. 2001). Male reproductive secretory tissues had divergent functions with the ejaculatory bulb contributing more differentially abundant proteins than the accessory glands, and the direction and severity of asymmetrical PMPZ isolation mirrors differential abundance. We also identified shared and species-specific GO enrichment of male reproductive proteins that influence reproductive processes, although whether PMPZ isolation more likely arises due to divergence in one or the other of these categories requires additional data. Democratisation of high throughput proteomics will facilitate understanding the evolution of male reproductive proteins, their influence on reproductive processes per se, and their contribution to reproductive isolation.

## ACKNOWLEDGEMENTS

Thanks to Priyanka Prajapati and Alexander Charles for assistance with proteomics data analysis, and Irem Sepil and the organisers and attendees of the Joint Wellcome Trust/EMBL-EBI Proteomics Bioinformatics course 2017 for discussion about analyses. Alison Wright, Daniela Palmer, Henry Barton and Toni Gossman provided helpful discussion and scripts for performing evolutionary rates analysis. We are grateful to Anneli Hoikkala for providing fly stocks, Michael G. Ritchie and Darren J. Parker for access to genomic and proteomic resources, and Roger Butlin for valuable feedback throughout the project. MDG was able to attend the Wellcome Trust/EMBL-EBI Proteomics Bioinformatics course thanks to a University of Sheffield Postgraduate Research Experience Programme (PREP) grant and was supported by the Adapting to the Challenges of a Changing Environment (ACCE) Doctoral Training Partnership grant NE/L002450/1, funded by the Natural Environment Research Council (NERC). Costs for proteomics was funded by a Royal Society Leverhulme Trust Senior Research Fellowship to RRS. CE acknowledges financial support from the Engineering and Physical Sciences Research Council, the ChELSI initiative (EP/E036252/1).

## AUTHOR CONTRIBUTIONS

RRS, TLK and CE conceived the study. RRS and MDG received funds for the work. MDG and CE collected the data. MDG and TLK analysed the data. MAR provided additional analysis tools. MDG and RRS wrote the manuscript with contributions from all authors. All authors agreed on the final version of the manuscript.

## DATA ACCESSIBILTY

Supplementary material, data and R code used to perform analyses can be found at: https://github.com/MartinGarlovsky/Dmon_ejaculate_proteomics. Additional data has been submitted to Dryad (https://doi.org/10.5061/dryad.pvmcvdnhw). The mass spectrometry proteomics data have been deposited to the ProteomeXchange Consortium (http://proteomecentral.proteomexchange.org) via the PRIDE partner repository (Vizcaíno et al. 2016) with the dataset identifier PXD019634.

## Notes

https://github.com/MartinGarlovsky/Dmon_ejaculate_proteomics

https://doi.org/10.5061/dryad.pvmcvdnhw

### Competing Interest Statement

The authors have declared no competing interest.

