## Supplementary methods and figures for "Seminal fluid protein divergence among populations exhibiting postmating prezygotic reproductive isolation" (from another version of this preprint: version 1, 2020-06-09)

June 2020

[1]Department of Animal and Plant Sciences, University of Sheffield, Sheffield S10 2TN, UK
[*]Current address: Department of Biology, Syracuse University, Syracuse NY 13244, USA
[2]Department of Chemical and Biological Engineering, The University of Sheffield, UK, Sheffield S10 2TN, UK
[3]Caris Life Sciences, Phoenix, USA
[4]Centre for Mechanisms of Evolution, The Biodesign Institute, Arizona State University, USA
[5]Department of Zoology, Stockholm University, Stockholm 106-91, Sweden

ORCiD: MDG: 0000-0002-3426-4341 CE: 0000-0003-4356-9216 MAR: 0000-0001-5522-5715 TLK: 0000-0002-4180-6583 RRS: 0000-0003-1852-1448

---

This markdown document provides supplementary material for Garlovsky, M. D., Caroline Evans, Mathew A. Rosenow, Timothy L. Karr and Rhonda R. Snook "Seminal fluid protein divergence among populations exhibiting postmating prezygotic reproductive isolation". Complete analyses and data are available on github. The mass spectrometry proteomics data have been deposited to the ProteomeXchange consortium via the PRIDE partner repository (Vizcaíno et al. 2016) with the dataset identifier PXD019634. The unfiltered *proteinGroups.txt* file exported from MaxQuant has been submitted to Dryad(Additional data has been submitted to Dryad.

### Supplementary methods

##### Tissue collection, protein extraction and purification

Twenty-one-day old males were anaesthetised with ether, the abdomen removed with insect pins, and placed in a drop of phosphate buffered saline (PBS). The whole reproductive tract (Fig. S1) was moved to a second drop of PBS, rinsed, the testes removed from the rest of the reproductive tract and discarded. The ejaculatory duct and bulb were separated from the accessory glands, intact where possible. Each tissue was washed in a third drop of PBS and placed in a LoBind Eppendorf tube containing 15µl lysis buffer (5% w/v sodium deoxycholate; 1% w/v sodium dodecyl sulphate) and protease inhibitor cocktail kept on ice. After reproductive tissues were harvested from 15 males the combined sample was freeze/thawed three times; placed on dry ice for 5 minutes, then placed in a water bath at 20ºC for 30 seconds and vortexed for 30 seconds. Each sample was then centrifuged at 20kG for 5 minutes at 4ºC, supernatant collected and placed in a new Eppendorf and stored at -80ºC until further processing. In total we collected three biological replicates for each tissue and each population. Biological replicate 1 consisted of 30 individuals per population (collected over two days) divided in to two equal volumes after tissue collection and processed separately. Two equal volumes of biological replicate 2 were loaded on to the mass spectrometer and biological replicate 3 was run in singlet, yielding a total of 5 LC-MS/MS runs for each tissue from each population (Fig. S1).

To quantify protein concentration to standardise loading volumes on to the mass spectrometer we ran 1µl of each sample with Laemmli buffer on an SDS-PAGE gel (Fig. S2) and performed densitometry with GelAnalyzer. Protein samples were processed with a HiPPR™ detergent removal kit (Thermo Fisher™, Catalogue number: 88305) and reduced by the addition of 2µl 50 mM tris(2-carboxyethyl)phosphine (TCEP) followed by incubation at 60ºC for 1 hour. Samples were allowed to cool then alkylated by addition of 1µl 200mM methyl methanethiosulfonate (MMTS) for 10 minutes at room temperature. Samples were then proteolytically digested using trypsin added at a ratio of 1:20 trypsin:protein. The digestion step was performed overnight at 37ºC, followed by drying to completion using vacuum centrifugation. All samples were then resuspended in 20µl 3% v/v acetonitrile, 0.1% v/v trifluoroacetic acid prior to analysis by liquid chromatography tandem mass spectrometry (LC-MS/MS).

##### LC-MS/MS data acquisition

LC-MS/MS was performed by nano-flow liquid chromatography (U3000 RSLCnano, Thermo Fisher™) coupled to a hybrid quadrupole-orbitrap mass spectrometer (QExactive HF, Thermo Scientific™). Peptides were separated on an Easy-Spray C18 column (75 µm x 50 cm) using a 2-step gradient from 97% solvent A (0.1% formic acid in water) to 10% solvent B (0.08% formic acid in 80% acetonitrile) over 5 min then 10% to 50% B over 75 min at 300 nL/min. A 105-minute MS data dependent acquisition (DDA) method was set up on the QExactive HF with settings: The full MS scan was from 375-1500 m/z acquired in the Orbitrap at a resolution of 120,000 in profile mode. Subsequent fragmentation was Top 10 in the HCD cell, with detection of ions in the Orbitrap using centroid mode, resolution 30,000. The following MS method parameters were used for MS1: Automatic Gain Control (AGC) target 1e6 with a maximum injection time (IT) of 60 ms and MS2: AGC target 1e5, IT of 60 ms and isolation window 2 Da. The intensity threshold was 3.3e4, normalized collision energy 27, charge exclusion was set to unassigned, 1, exclude isotopes was on, apex trigger deactivated. The peptide match was set to preferred with dynamic exclusion of 20 seconds.

##### LC-MS/MS data processing

We performed quantitative proteomic analysis for label free quantification using the MaxLFQ algorithm to generate relative peptide and protein intensity information (Cox et al. 2014) in MaxQuant (Tyanova et al. 2016). We set carbamidomethyl as a fixed modification and acetylation of N-terminus of proteins and oxidation of methionine as variable modifications. For identification a minimum peptide length of 7 was required and a minimum ratio count of two for quantification. False discovery rate was set at 0.01 calculated using a reverse decoy database which we subsequently removed along with contaminants. For protein identification we matched mass spectra to the D. montana predicted proteome. Detailed description of the proteome construction can be found in (Parker et al. 2018). Briefly, the proteome was generated using gene predictions from the Maker2 pipeline (Holt and Yandell 2011) reciprocally blasted against D. virilis proteins (Parker et al. 2018).

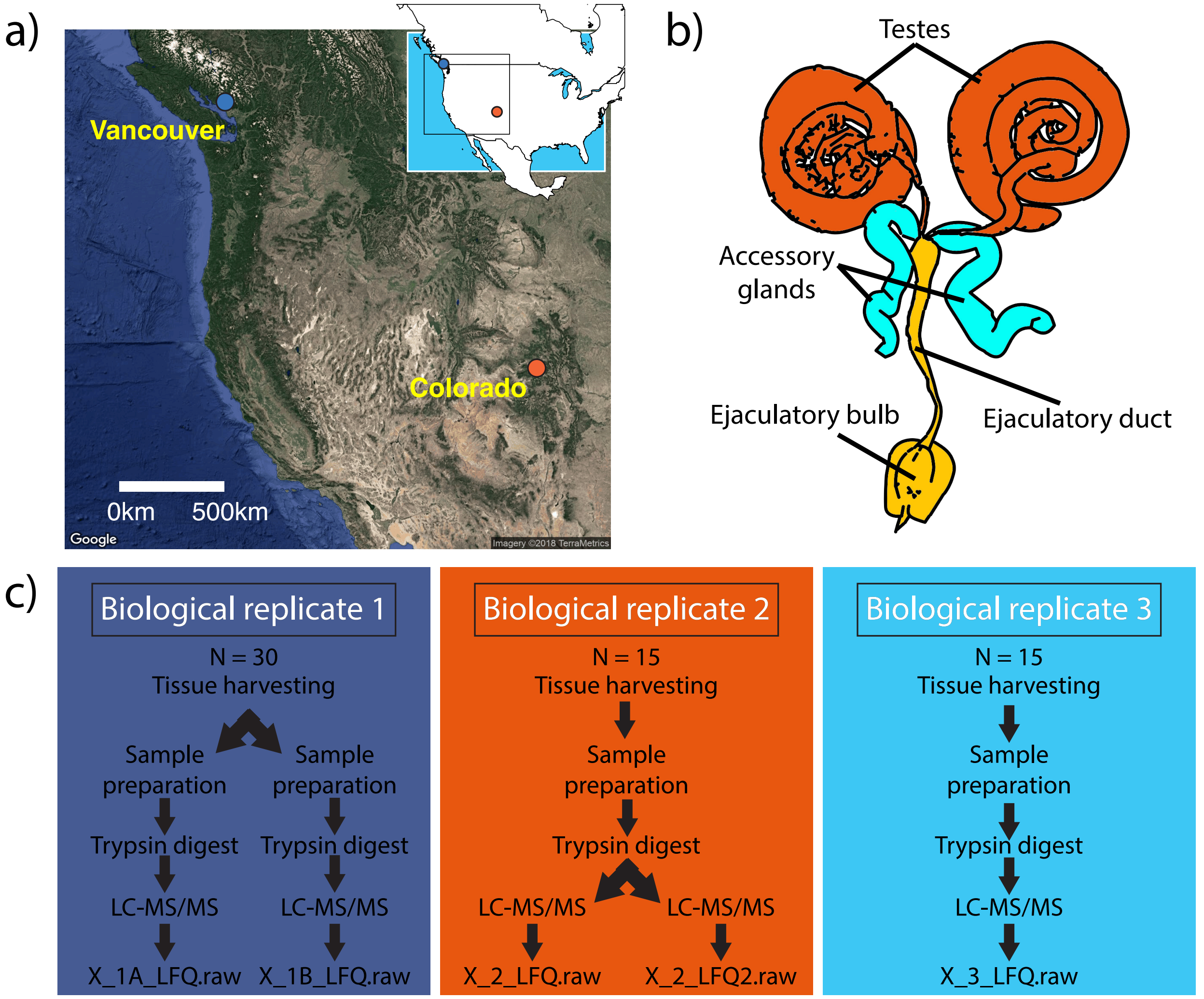


**Figure S1:** a) Collection locations of *Drosophila montana* populations. b) Male *D. montana* reproductive tract highlighting paired accessory glands (turquoise) and ejaculatory duct and bulb (yellow). Image modified from FlyBase.org after Patterson (1943). c) LC-MS/MS data acquisition experimental design. N = number of male reproductive tracts pooled in each replicate. "X_1A_LFQ.raw" etc. refer to nomenclature used for naming raw mass spectra data. "X" refers to population (Colorado [C] or Vancouver [V]) followed by tissue type (accessory glands [AG] or ejaculatory ducts and bulbs [EB]); "X_1A/1B" = sample processing technical replicates of biological replicate 1; "X_2_1/2" = LC-MS/MS technical replicates of biological replicate 2; "X_3" = biological replicate 3, ran in singlet.

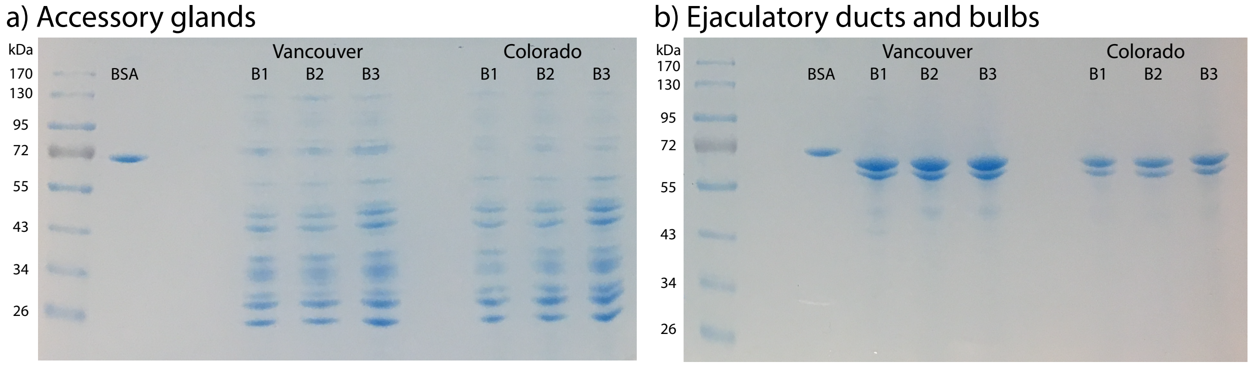


**Figure S2:** SDS-PAGE gel profiles of a) accessory glands and b) ejaculatory ducts and bulbs. One microlitre of each tissue sample was diluted in Laemmli buffer and loaded plus 1µL of Bovine Serum Albumin (BSA) standard (0.15µg/µl). B1-3; Biological replicates 1-3. Images have been cropped. Molecular weight ladder: Fisher BioReagents™ EZ-Run™ Prestained Rec Protein Ladder, range 10 -170 kDa Cat no. BP3603500.

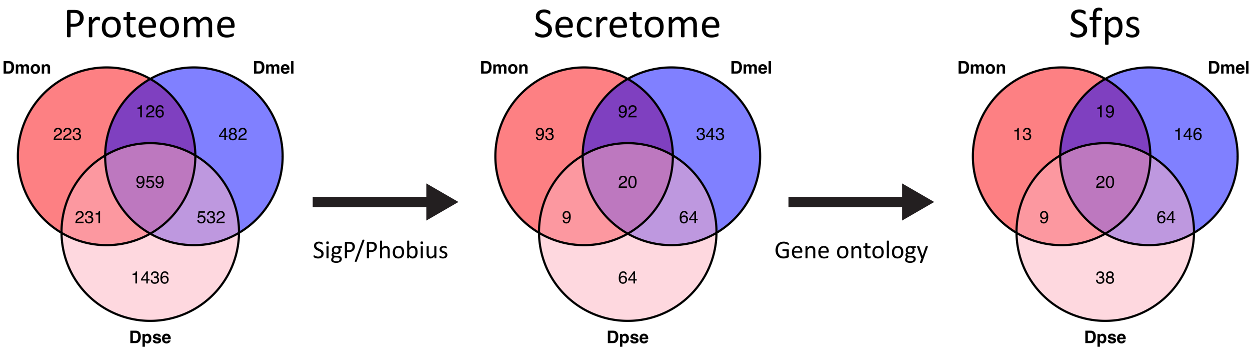


**Figure S3:** Construction of secretomes and seminal fluid protein (Sfp) lists. We retrieved FBgns for *D. melanogaster* genes identified by Sepil et al. (2019) and *D. melanogaster* orthologs for *D. pseudoobscura* genes identified by Karr et al. (2019) and downloaded the corresponding canonical protein sequences from uniprot.com. For *D. montana* we retrieved protein sequences from Parker et al. (2018) for proteins we identified in our analysis. We submitted protein sequences for each species to *SignalP* (Petersen et al. 2011) and *Phobius* (Käll et al. 2004) and combined the resulting lists of proteins containing a signal peptide to generate a list of secretome proteins for each species. For *D. montana* we converted the corresponding *D. virilis* FBgns for each protein to *D. melanogaster* orthologs via FlyBase.org (for 215/245, 88%). To identify Sfps for each species we submitted secretome protein lists to FlyBase.org to retrieve genes with GO terms containing "extracelluar".

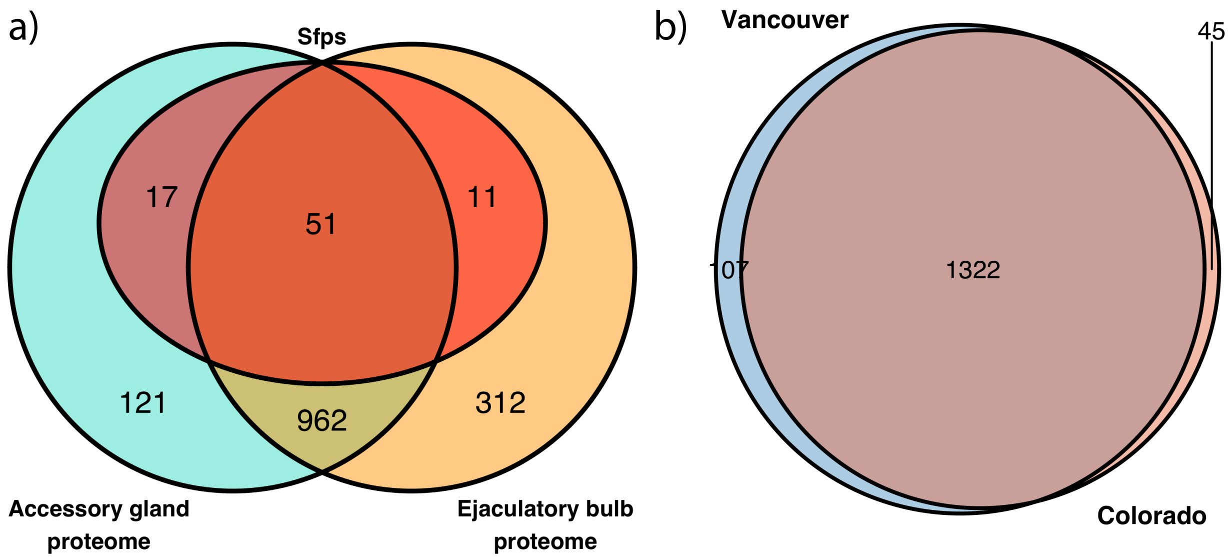


**Figure S4:** Number of proteins identified in each tissue a) and each population b). a) We found 1474 *D. montana* proteins identified by at least two unique peptides, including 79 seminal fluid proteins (Sfps). The list of Sfps was compiled by combining the 55 proteins identified by 2 or more unique peptides with a predicted signal peptide sequence and extracellular annotations via FlyBase.org plus 38 orthologs of *D. melanogaster* Sfps identified by converting *D. virilis* FBgns to *D. melanogaster* orthologs via FlyBase.org. b) The majority of the 1474 proteins were identified in both populations (1322, 90%), while 45, and 107 were only identified in Colorado, and Vancouver, respectively.

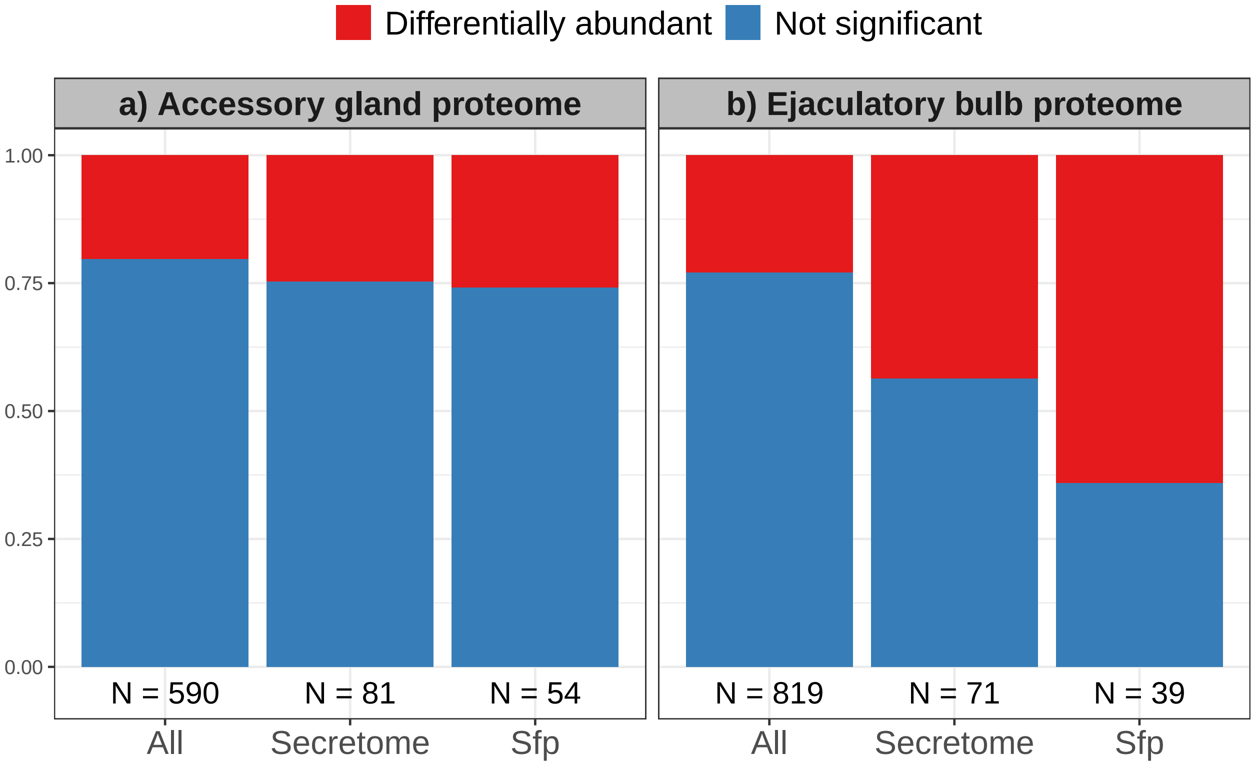


**Figure S5.** Representation of secretome and seminal fluid proteins (Sfps) in the set of differentially abundant proteins between populations. Secretome proteins and Sfps were equally represented in the set of differentially abundant proteins between Colorado and Vancouver in the accessory gland proteome (Chi-squared test, $\chi^2 = 1.57$, df = 2, p = 0.456) but were overrepresented in the differentially abundant proteins in the ejaculatory bulb proteome ($\chi^2 = 44.56$, df = 2, p < 0.001). N = sample sizes.

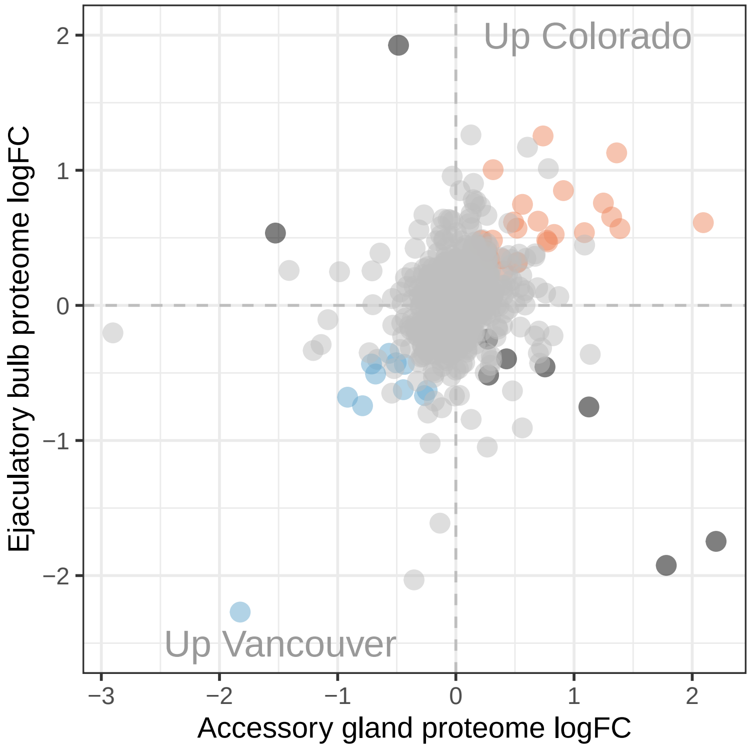


**Figure S6.** Differentially abundant proteins between Colorado and Vancouver found in both the accessory gland proteome and ejaculatory bulb proteome. Proteins showing differential abundance between populations in both tissues showing significant and concordant log2-fold-change abundance are highlighted for Colorado (red, n = 25), Vancouver (blue, n = 11), or discordant (black, n = 9).

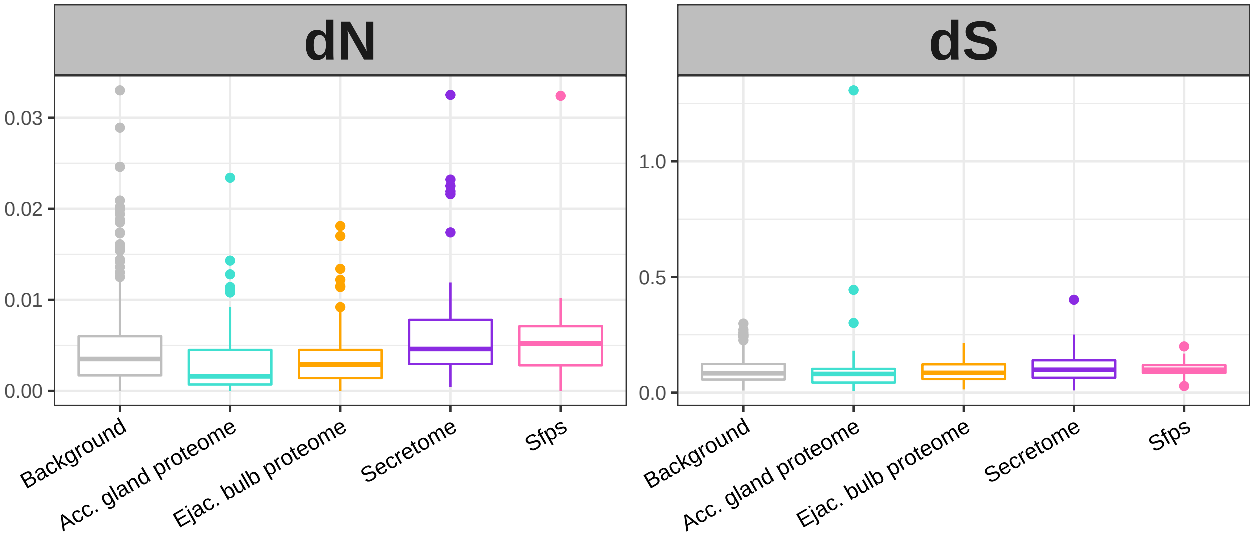


**Figure S7.** Estimates of non-synonymous (dN; left panel) and synonymous (dS; right panel) substitution rates for *D. montana* genes. Genes are categorised based on showing equal protein abundance in the accessory gland proteome or ejaculatory bulb proteome ('background'; grey), higher abundance in the accessory gland proteome (turquoise), ejaculatory bulb proteome (orange), or found in the secretome (purple) or Sfps (pink).
